# Integrated population clustering and genomic epidemiology with PopPIPE

**DOI:** 10.1101/2024.12.05.626978

**Authors:** Martin P. McHugh, Samuel T. Horsfield, Johanna von Wachsmann, Jacqueline Toussaint, Kerry A. Pettigrew, Elzbieta Czarniak, Thomas J. Evans, Alistair Leanord, Luke Tysall, Stephen H. Gillespie, Kate E. Templeton, Matthew T. G. Holden, Nicholas J. Croucher, John A. Lees

**Affiliations:** Medical Microbiology, Department of Laboratory Medicine, Royal Infirmary of Edinburgh, Edinburgh, UK; Division of Infection and Global Health, University of St Andrews, St Andrews, United Kingdom; European Molecular Biology Laboratory, European Bioinformatics Institute EMBL-EBI, Hinxton, United Kingdom; School of Infection and Immunity, University of Glasgow, Glasgow, United Kingdom; Scottish Microbiology Reference Laboratories, Glasgow Royal Infirmary, Glasgow, United Kingdom; MRC Centre for Global Infectious Disease Analysis, School of Public Health, Imperial College London, London, United Kingdom

**Keywords:** Genomic epidemiology, clustering, transmission, pipelines

## Abstract

Genetic distances between bacterial DNA sequences can be used to cluster populations into closely related subpopulations, and as an additional source of information when detecting possible transmission events. Due to their variable gene content and order, reference-free methods offer more sensitive detection of genetic differences, especially among closely related samples found in outbreaks. However, across longer genetic distances, frequent recombination can make calculation and interpretation of these differences more challenging, requiring significant bioinformatic expertise and manual intervention during the analysis process. Here we present a **Pop**ulation analysis **PIPE**line (PopPIPE) which combines rapid reference-free genome analysis methods to analyse bacterial genomes across these two scales, splitting whole populations into subclusters and detecting plausible transmission events within closely related clusters. We use k-mer sketching to split populations into strains, followed by split k-mer analysis and recombination removal to create alignments and subclusters within these strains. We first show that this approach creates high quality subclusters on a population-wide dataset of *Streptococcus pneumoniae*. When applied to nosocomial vancomycin resistant *Enterococcus faecium* samples, PopPIPE finds transmission clusters which are more epidemiologically plausible than core genome or MLST-based approaches. Our pipeline is rapid and reproducible, creates interactive visualisations, and can easily be reconfigured and re-run on new datasets. Therefore PopPIPE provides a user-friendly pipeline for analyses spanning species-wide clustering to outbreak investigations.

**Impact statement:** As time passes, bacterial genomes accumulate small changes in their sequence due to mutations, or larger changes in their content due to horizontal gene transfer. Using their genome sequences, it is possible to use phylogenetics to work out the most likely order in which these changes happened, and how long they took to happen. Then, one can estimate the time that separates any two bacterial samples – if it is short then they may have been directly transmitted or acquired from the same source; but if it is long they must have been acquired separately. This information can be used to determine transmission chains, in conjunction with dates and locations of infections. Understanding transmission chains enables targeted infection control measures. However, correctly calculating the genetic evidence for transmission is made difficult by correctly distinguishing different types of sequence changes, dealing with large amounts of genome data, and the need to use multiple complex bioinformatic tools. We addressed this gap by creating a computational workflow, PopPIPE, which automates the process of detecting possible transmissions using genome sequences. PopPIPE applies state-of-the-art tools and is fast and easy to run – making this technology will be available to a wider audience of researchers.

**Data summary:** The code for this pipeline is available at https://github.com/bacpop/PopPIPE and as a docker image https://hub.docker.com/r/poppunk/poppipe. Raw sequencing reads for *Enterococcus faecium* isolates have been deposited at the NCBI under BioProject accession number PRJNA997588.

## Introduction

Routine genomic surveillance and targeted sequencing of bacterial pathogens have become central tenets of infectious disease epidemiology. Analysis of genomes from bacterial pathogens, at its simplest, will give information on the species and strain (also called cluster, sequence type, lineage or clade) of the sequenced isolate. Typically these are two compatible hierarchical definitions, often made at 95% and 99.5% sequence identity respectively [1–3]. Genetically determined species largely agree with traditional prokaryotic taxonomy and determines microbiologically and clinically important differences between infectious agents [4]. Strain designations sometimes have useful correlation with phenotypes (e.g. virulence, resistance) due to inherited genetic elements within the clades they represent.

Strain designations are powerful at *ruling out* recent transmission between samples, as the genetic distance between two strains frequently represent hundreds or thousands of years of evolution [5]. A further level of sub-clustering within strains, which we refer to as lineages, can be useful for making this ruling out more sensitive (without loss of precision). However, genetic distances alone are less powerful at *ruling in* transmission [6], and for this purpose are best combined with epidemiological data on sampling time [7] and place [8]. To bring these insights from complex models into epidemiological practice also requires judicious use of suitable interactive visualisations [9, 10].

Integrated genomic and epidemiological analysis and visualisation of multi-strain bacterial populations are typically complicated due to the fact that across the entire species a vertical evolutionary history cannot be accurately reconstructed from contemporary genomes alone [11]. If this isn’t accounted for, genetic branch lengths are likely to be overestimated, causing significant errors in sub-clustering and transmission analyses [12].

Furthermore, selecting correct and efficient methods from a wide array of possibilities, connecting them together and making this reproducible is a challenge. For epidemiological analysis using genomes, particularly if this analysis is going to be used to inform interventions or policy changes, it is critical that results are reliable and reproducible. In some cases where external validation of pipelines is required for use in public health, their constituent programs and parameters must remain stable so their evaluation performance on validation datasets is unchanged [13]. Even for experienced bioinformaticians, best practices from multiple pieces of software can be hard to follow or adapt exactly to the use case in hand. These issues can be helped by pipeline managers such as Snakemake and Nextflow [14, 15]. Using these managers, pipelines can run with set programs and versions, and transparently set configuration which can be applied across different datasets.

We therefore designed PopPIPE (**Pop**ulation analysis **PIPE**line) to automate the complex orchestration of running many bioinformatic tools for sub-clustering and transmission analysis across bacterial populations with many stains. By using sensible, but easily adjustable, defaults and efficient choices of bioinformatic software tools, the pipeline is rapid to run, requires minimal user input, and produces reliable and reproducible genomic inferences. Our pipeline has three modes: sub-clustering, visualisation and transmission detection. We demonstrate its operation on a population-wide dataset of *Streptococcus pneumoniae*, and a hospital outbreak of vancomycin resistant *Enterococcus faecium*.

## Theory and implementation

Phylogenetics and genomic-based transmission detection rely on an accurate clonal frame where vertical evolution can be inferred as a series of sequential events along a tree. If recombination events are not removed prior to these steps, branch lengths will be overestimated [12] and transmission events/clusters missed. Methods exist to remove these recombination events (for example gubbins [16], ClonalFrameML [17]), but are only applicable within populations of limited diversity sharing a recent common ancestor.

Many bacterial species have populations which maintain a genetic structure of different strains (also called sequence types). Analogous to an average nucleotide identity of 95% separating most species, an average nucleotide identity of around 99% separates most strains [18–20]. However, it is often challenging to reconstruct any valid vertical evolutionary history across these strains due to frequent and repeated overwriting of sequences resulting from homologous recombination and the movement of mobile genetic elements [11]. Although gene-based methods such as FastGEAR [21] can detect ancestral recombination events, these cannot capture the whole-genome resolution needed to accurately distinguish closely related samples in an outbreak setting.

These are complex evolutionary considerations that may not always be easy to address or within the scope of a genomic epidemiology solution. We therefore present a pragmatic and general solution to this issue: a population is first subdivided into genetic clusters *between which* transmission is completely implausible, and *within which* a recent common ancestor exists. Then, in every such cluster recombination can be detected and removed, leaving a clonal frame from which phylogenetics and transmission detection can be performed. Specifically, within each cluster we aim to produce the following useful components for genomic epidemiology analyses:

- Genetic subclusters, defined hierarchically with multiple nested levels of further subclusters.
- A maximum-likelihood phylogeny from whole-genome alignment.
- A recombination removed, dated phylogeny.
- Inferred transmission events based on this phylogeny.
- Interactive visualisation of the genetic relationships and subclusters for the whole dataset.

We refer to these subclusters as lineages, due to analogous usage in the non-multi-strain *Mycobacterium tuberculosis* [22] and viral pathogens [23].

We implemented this process in PopPIPE (v1.1.0), a Snakemake (v7.8.5) pipeline [14]. The pipeline can therefore continue failed or paused analysis from where it left off, rerun the same analysis on new data completely consistently with previous runs, and automatically take advantage of multicore systems or common job submission systems. Our pipeline is designed to be run following a PopPUNK strain-typing analysis [2], which is first used to split a population into strains. The data needed as input is:

- A user’s self-made PopPUNK database, or an existing PopPUNK database if stable strain nomenclature is desired (https://www.bacpop.org/poppunk/).
- Assemblies (if available, preferred for speed) or Illumina sequence reads for all samples included.
- If transmission is to be inferred, a file listing the dates all samples were taken.
- A config file, listing file paths of these inputs, and defining parameters for tools in the pipeline.

The steps of the default pipeline which makes subclusters of the data is as follows (Figure 1):

**Figure 1:**
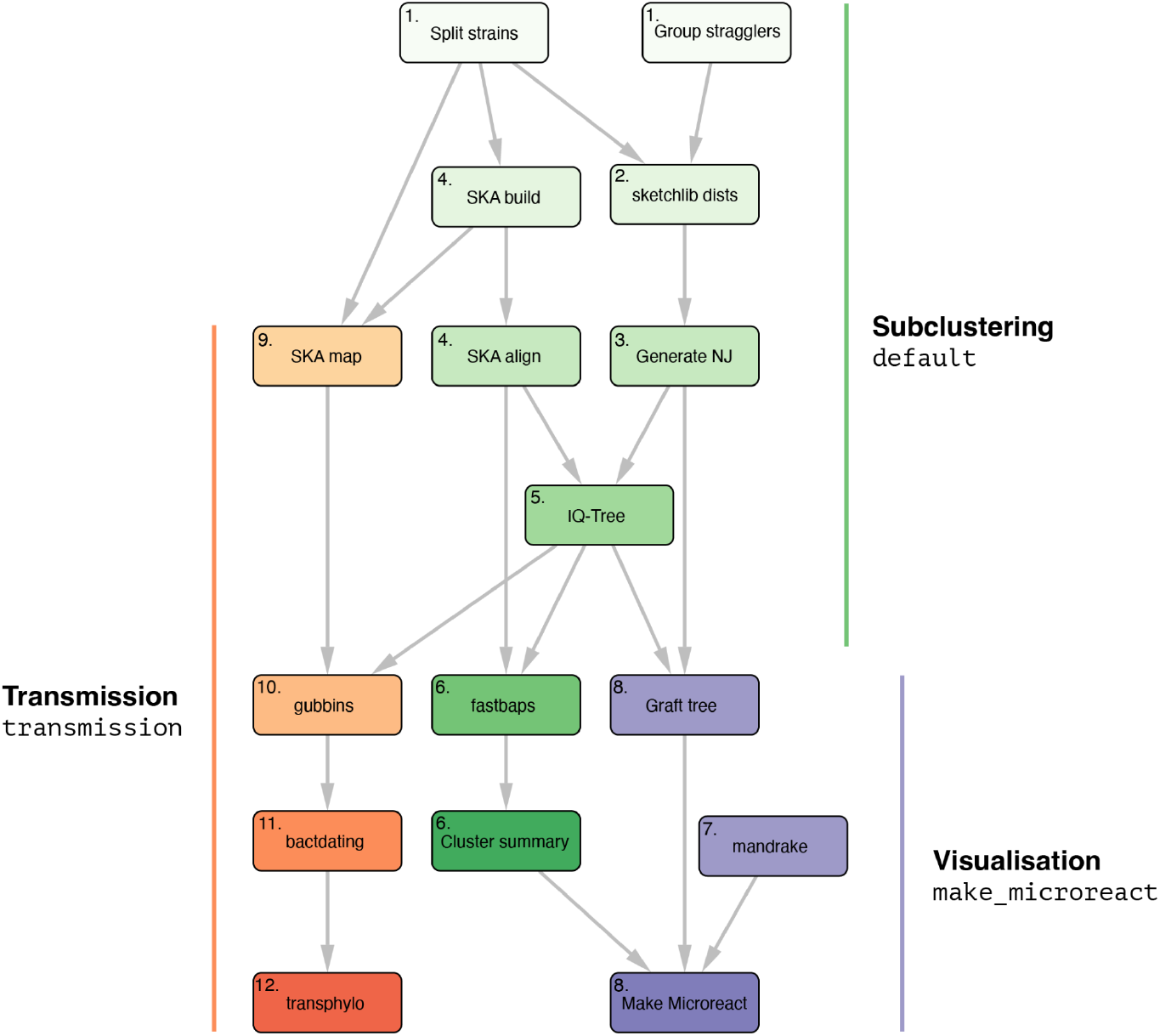
Graph representing steps in PopPIPE. Nodes are snakemake rules, usually running a piece of software, edges are dependencies where a step will only start running when all its dependencies have completed. Nodes are annotated with the step number where they are fully described in the text. Green nodes are run for the `default` target of generating sub-clusters. Purple nodes are run when a visualisation is being made with the `make_microreact` target. Orange nodes are run when transmission detection is run with the `transmission` target.

1. Partition assembly/read files into their `PopPUNK` (v2.7.0) [2] strains. Steps 2-6 are then carried out in parallel across each strain.
2. Use `pp-sketchlib` (v2.1.3) to calculate core and accessory distances within each strain, using the same parameters as the PopPUNK database.
3. Use the core distances with `rapidnj` (v2.3.2) [24] to make a neighbour-joining tree.
4. Use SKA2 (v0.3.9) [20] to generate reference-free alignments using `ska align`.
5. Use `IQ-TREE` (v2.0.3) [25] to generate a maximum-likelihood phylogeny using this alignment, and using the neighbour-joining tree as a starting point.
6. Use `fastbaps` (v1.0.5) [26] to generate subclusters/lineages, in the mode where subclusters are necessarily partitions of the phylogeny (increasing speed and eliminating polyphyletic clusters).

To make an interactive visualisation of the subclusters in microreact [27] two further steps are run:

7. Use `mandrake` (v1.2.1) [28] to create an embedding visualisation of the accessory distances.
8. Create an overall tree by grafting the maximum likelihood trees for subclusters to their matching nodes, rescaling branch lengths to match the neighbour-joining tree, and midpoint rooting maximum-likelihood trees. We note that this tree will not maximise the phylogenetic likelihood – it is only intended as a convenient visualisation of the entire dataset, which is compatible with existing tools.

This creates a .microreact file as output, and if an API key is provided will automatically create a permanent URL hosting the visualisation.

To infer transmission trees, four further steps are run:

9. SKA2 generates alignments with `ska map` and a reference genome from the strain so SNPs form correct windows on the physical chromosome.
10. Use `gubbins` (v3.1.0) [16] to remove recombination.
11. Use `bactdating` (v1.1) [29] with sampling times to infer timed trees from recombination-purged phylogenies.
12. Use `transphylo` (v1.4.8) [7] to infer possible transmission events on these timed trees.

## Sub-clustering and visualisation of *S. pneumoniae*

We demonstrated PopPIPE on 616 previously published *S. pneumoniae* genomes [19, 30]. PopPUNK database construction and clustering took two minutes using four threads. Using PopPIPE to create a visualisation with a phylogeny and subclusters took an additional 28 minutes using four threads. The majority of this time was spent on maximum likelihood tree inference – 50% of the total was spent on inferring the tree of largest strain (98 samples). Of the 62 inferred strains, 28 had a sufficient number of members to subcluster (>=6), producing 101 first level subclusters and 157 second level subclusters (Figure 2). Running the transmission pipeline took an additional 52 minutes, the majority of the time being spent on recombination removal.

**Figure 2:**
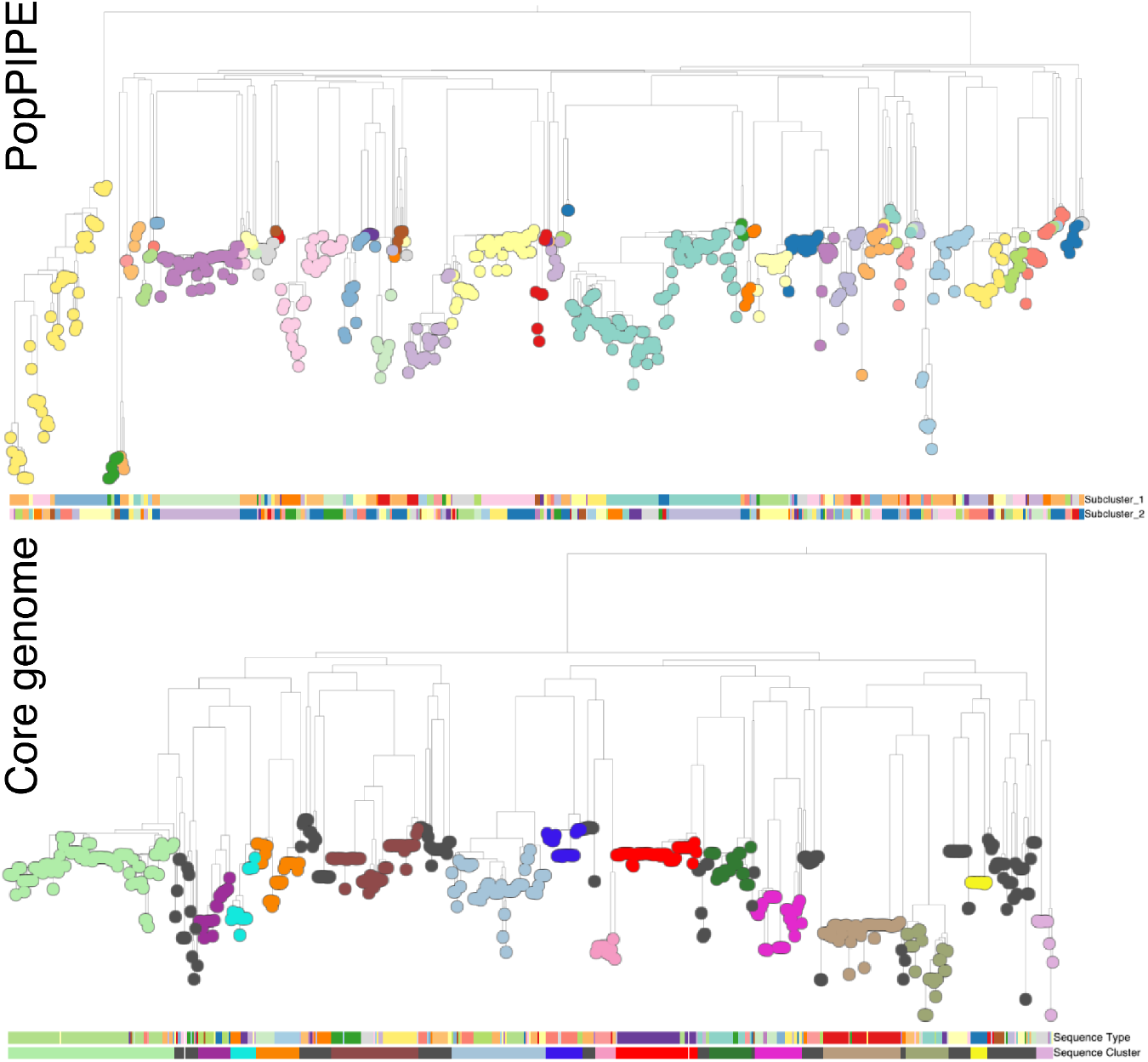
Visualisation results from the PopPIPE sub-cluster analysis (top) compared to the original core genome and sequence-type analysis (bottom).

The results of this pipeline run can also be found online (https://microreact.org/project/sparc-poppipe) and in Supplementary Figure 1. We compared this to the original analysis of this data (also available on microreact https://microreact.org/project/NJwviE7F), which used a concatenated core gene alignment followed by maximum likelihood tree construction [19], Qualitatively similar results were found for tree topology and clusters (Figure 2). A comparison of the tree topology in phylo.io [31] showed differences in the ancestral branches of the tree topology, as expected due to unresolvable recombination events, but generally close concordance near the tips within individual strains (Supplementary Figures 2-3). The rooting and topology used in PopPIPE is not optimised, so gives a worse tree. However, by construction, the subclusters/lineages necessarily matched the tree and its resolution, whereas sequence types offered inconsistent resolution across the tree as observed in [32].

As a more quantitative comparison, we also compared the topology of all of the subtrees for the 39 strains with more than three samples in the PopPIPE data using the Kendall-Colijn tree metric [33]. Ten were identical, 24 had only minor topology changes (distance < 20 and confirmed visually as in [34]). The remaining five strains were the larger strains, the most divergent of which was strain 4 (GPSC59; ST558; serotype 35B [35]) with a Kendall-Colijn distance of 177. This subtree has long terminal branches, a signal typical of unremoved recombination. Using the final subtree from PopPIPE, after recombination has been removed, results in a tree which is closer to the original, with a Kendall-Colijn distance of 95. Overall, this shows good phylogenetic concordance from this rapid pipeline when compared to a high-quality manual analysis.

## Transmission detection in vancomycin resistant *Enterococcus faecium*

Several recent studies have also shown the utility of alignment-free clustering for investigation of *E. faecium* transmission [36–39]. Higgs et al [36] report that SKA was more suitable than reference based mapping to infer patient-to-patient transmission of *E. faecium*, and recommend the use of SKA for this task going forward. Maechler et al [37] compared clustering with core genome MLST (cgMLST), core genome SNPs, and SKA with similar findings to those reported here – SKA generated more clusters with fewer genomes per cluster, but higher agreement with epidemiological data than both cgMLST and core SNPs. Both Maechler et al [37] and Rath et al [39] show SKA is suitable for *E. faecium* clustering in nosocomial settings and helps to rule in and rule out possible transmission links in combination with epidemiological data.

We therefore demonstrated the use of SKA within PopPIPE to subcluster a local outbreak dataset. Vancomycin resistant *Enterococcus faecium* (VREfm) readily spreads in healthcare settings leading to challenging outbreaks. Whole genome sequencing (WGS) is increasingly used to identify putative transmission and target infection control measures. Short-read Illumina WGS was performed on 87 VREfm isolates from 84 patients included in an outbreak investigation across six orthopaedic inpatient wards. Adapters and low quality regions were removed with Trimmomatic (v0.39) [40] then the trimmed reads were assembled with SPAdes (v3.15.5) [41] using the --isolate flag. PopPUNK was used to create a database and subsequently cluster the assemblies. PopPIPE was then used to subcluster and infer transmission trees with transphylo. PopPIPE clustering was compared with reference-based core genome SNP clustering. Briefly, trimmed reads were mapped to the Aus0004 reference genome (accession CP003351) and mobile genetic elements masked using Snippy (v4.6.0) (https://github.com/tseemann/snippy), recombination was identified with Gubbins [16] and masked from the alignment, SNPs were extracted using SNP-sites (v2.4.0) [42], and pairwise SNP counts calculated. Finally, core genome SNP clusters were identified with a threshold of ≤3 SNPs, as this was previously identified as the maximum diversity seen within individual patients [43]. Patient stay metadata was retrieved from electronic health records.

PopPIPE assigned 77 (88.6%) genomes into 20 clusters, while core genome SNPs grouped 74 (85.1%) genomes into 14 clusters. PopPIPE and core genome SNP grouping agreed in 68 (78.2%) cases. Within PopPIPE clusters, 30% of patient pairs were on the same ward at the same time and 3% had no epidemiological link, compared to 26% and 5% in core genome SNP clusters (Table 1). VREfm PopPIPE cluster 1-4-5 contained nine patients and was further examined as an example of transmission inference (Figure 3, Supplementary Figure 4). The transmission tree suggested P1 transmitted to P4, both patients shared time and place in the hospital. P1 was also inferred to have transmitted to P8, who was on the same ward but at a different time; P34, who was on a different ward at the same time, and P30, who was epidemiologically unlinked. P8 was inferred to have transmitted to P19 and P21, all of whom were patients on the same ward at the same time. P4 was inferred to have transmitted to P16, these patients had no time or ward overlap in the hospital. P57 tested positive on screening at a pre-admission clinic, with no contact with the hospital in the preceding 6 months. This case was not linked to direct transmission in the transmission tree which suggests pre-existing carriage from an unidentified source. TransPhylo was useful in ruling out possible direct transmission (P57) and for indicating likely linked patients (P1/P4 and P8/P19/P21) but for other cases the inference should be examined with epidemiological data before considering direct transmission links. This reflects the challenges in inferring direct transmission, including the uncertain serial interval for *E. faecium*, long asymptomatic carriage, potential hospital environmental reservoirs, and unsampled hosts.

**Table 1:** Patient epidemiological linkage within VREfm genomic clusters.

| **Epidemiological linkage** | **PopPIPE [n=159]**<br>**% (95% CI)** | **Core Genome**<br>**SNP [n=228]**<br>**% (95% CI)** |
| --- | --- | --- |
| Same ward, same time | 30.2 (23.1-37.3) | 25.9 (20.2-31.6) |
| Same ward, stay 1-28<br>days apart | 12.6 (7.4-17.8) | 14.1 (9.6-18.6) |
| Different ward, same time | 26.5 (19.6-33.4) | 22.0 (16.6-27.4) |
| Different ward, stay 1-28 days apart | 27.7 (20.7-34.7) | 32.9 (26.8-39.0) |
| No link | 3.2 (0.5-5.9) | 5.3 (2.4-8.2) |

CI, confidence interval; n = comparison pairs; SNP, single nucleotide polymorphism

**Figure 3:**
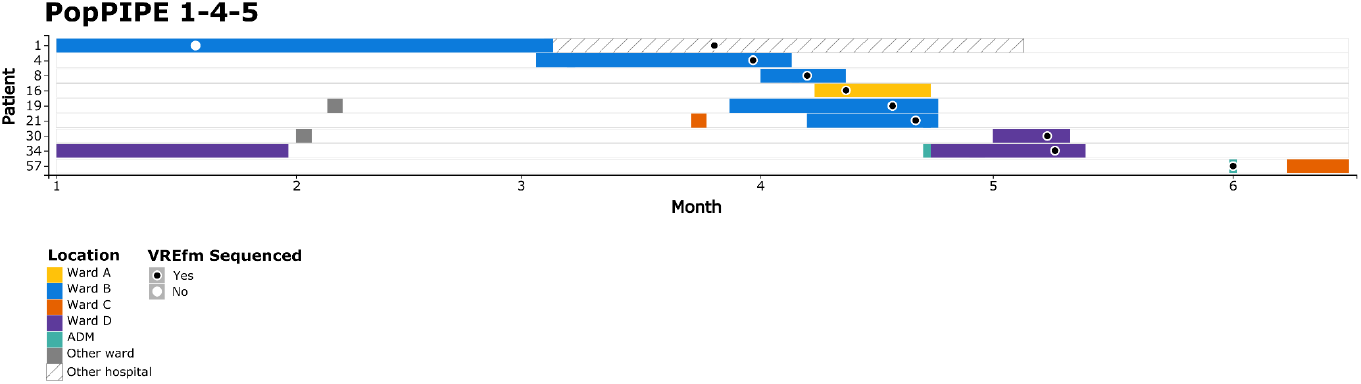
Patient timelines for PopPIPE cluster 1-4-5. Each patient is represented by a row, with month on the x-axis. Hospital visits are plotted as coloured boxes, coloured by the ward, isolates are represented by circles and coloured based on whether the isolate was sequenced or not. ADM, pre-admission clinic. Figure generated with HAIviz (v0.3) [44].

## Discussion

Recent computational methods use reference-free approaches to compare bacterial genomes more quickly and sensitively than traditional reference-based approaches. Sketching methods such as PopPUNK use variable length k-mers to rapidly calculate genetic distances across entire species, and produce epidemiological clusters that divide populations into strains (usually separated by at least decades of evolution). SKA is a variant calling method which uses split k-mers to rapidly find specific variants within these strains. Combining sketching methods to split a population into first-level clusters, with fast variant calling within these clusters to obtain further subclusters, is an appealing approach to work with large sets of bacterial genomes. This combination of methods breaks analysis into manageable pieces which correspond to subpopulations where the vertical inheritance can be reconstructed with a degree of accuracy. Such methods are particularly needed in an era where millions of publicly available genomes are available to give context to population analyses [45], and where thousands of new genomes are sequenced every day [46].

By building on these underlying reference-free methods, PopPIPE represents a fast and reproducible method for the fundamental steps of population genetics using sets of genomes. What was previously the analysis underpinning an entire research project [19] can be completed automatically in under an hour, with the option to rerun or restart the same analysis with different parameters, or be reproduced exactly by other users.

Genetic distances are also a potentially powerful addition to infectious disease epidemiology methods, supplementing traditional ‘line lists’ with case times and location with an additional data source. Though genetic distances alone cannot typically resolve who-infected-whom [47], the combination of plausible evolutionary time, sampling time and physical proximity is a sensitive and specific way to detect transmission chains [8]. To obtain the best resolution on genetic distances in bacterial pathogens requires a reference-free and pangenome-wide approach [48], as traditional reference-based read-mapping methods will suffer from biases leading to decreased sensitivity and specificity [49]. Although SKA, the underlying reference-free method, is a fast and sensitive method to determine between sample genetic distances, it does not infer likely transmissions directly. Additionally, to achieve precision across a population requires careful prior clustering and recombination removal. We showed here that automating these steps in PopPIPE led to more precise and epidemiologically plausible transmission clusters than a core genome analysis.

We designed PopPIPE as an accessible route for researchers from all bioinformatic backgrounds to achieve high-quality SKA-based sub-clustering and transmission analysis. PopPIPE brings together a multitude of modern tools, initially designed to be run individually, together, enabling reference-free epidemiology. PopPIPE is therefore well placed for routine outbreak investigations given its speed, built-in visualisation options, and ease of repeating reproducible analyses with new data.

## Author statements

### Author contributions

Conceptualisation: MPM, NJC, JAL. Data curation: MPM, KAP, EC, DI, LT, NJC. Formal analysis: MPM, JAL. Funding acquisition: TJE, AL, SHG, MTGH, NJC, JAL. Methodology: JAL, JvW. Software: STH, JvW, JT, JAL. Resources: MPM, JAL. Supervision: SHG, KET, MTGH, NJC, JAL. Validation: JT, NJC. Visualization: MPM, JAL. Writing – original draft: MPM, JAL. Writing – review & editing: all authors.

### Conflicts of interest

The author(s) declare that there are no conflicts of interest.

### Funding information

S.T.H., J.v.W.., J.T. and J.A.L. were supported by the European Molecular Biology Laboratory. S.T.H. and J.A.L. were additionally supported by BBSRC (reference BB/Y513805/1). N.J.C. was supported by the UK Medical Research Council and Department for International Development (grants MR/R015600/1 and MR/T016434/1). N.J.C. acknowledges funding from the MRC Centre for Global Infectious Disease Analysis (reference MR/X020258/1 and MR/T016434/1), funded by the UK Medical Research Council (MRC). This UK funded award is carried out in the frame of the Global Health EDCTP3 Joint Undertaking. M.P.M., K.A.P., T.J.E., A.L., S.H.G, and M.T.G.H were supported by the Chief Scientist Office (Scotland) (grant number SIRN/10). Analysis of *E. faecium* data was performed on the Crop Diversity Bioinformatics HPC hosted by the James Hutton Institute and funded by BBSRC (reference BB/S019669/1).

### Ethical approval

Access to *E. faecium* isolates as excess diagnostic material was approved by the NHS Scotland BioRepository Network (reference TR000126). Access to enhanced patient metadata was approved by the NHS Lothian Caldicott Guardian (reference 1690).

## Acknowledgements

We would like to thank Dr Donald Inverarity for assistance with Enterococcus faecium outbreak data.

## Notes

https://github.com/bacpop/PopPIPE

### Competing Interest Statement

The authors have declared no competing interest.

