## Supplementary material for "Integrated population clustering and genomic epidemiology with PopPIPE"

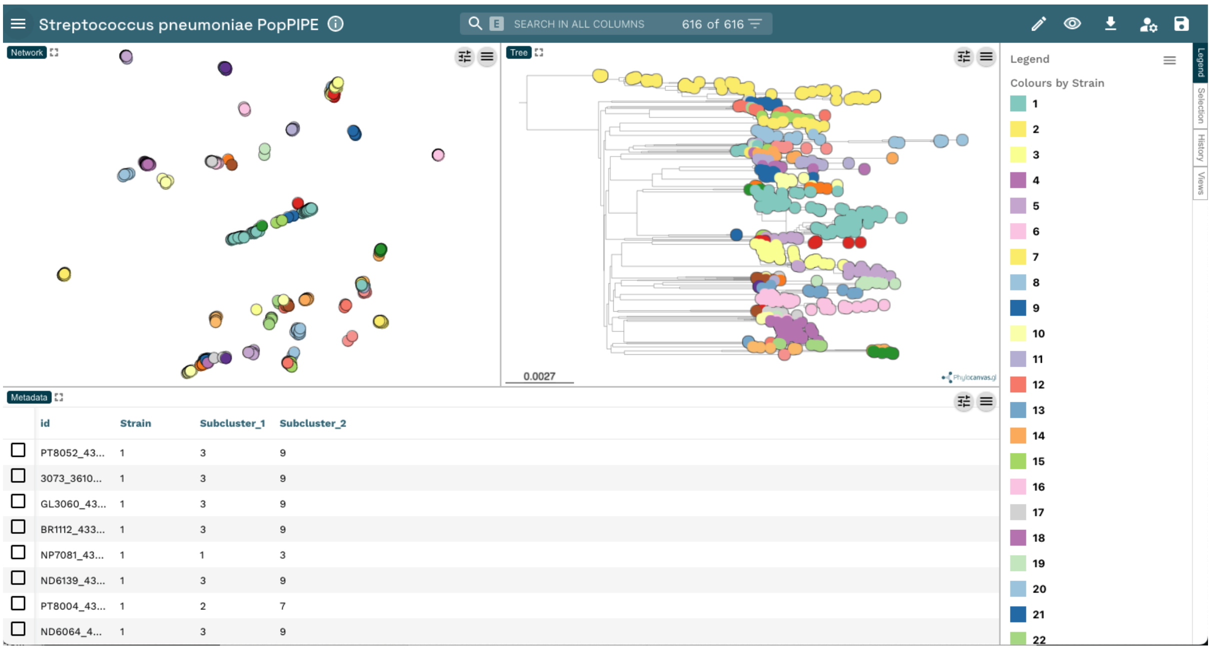


**Supplementary figure 1:** Full visualisation of the PopPIPE run from figure 5. Also available online: https://microreact.org/project/sparc-poppipe

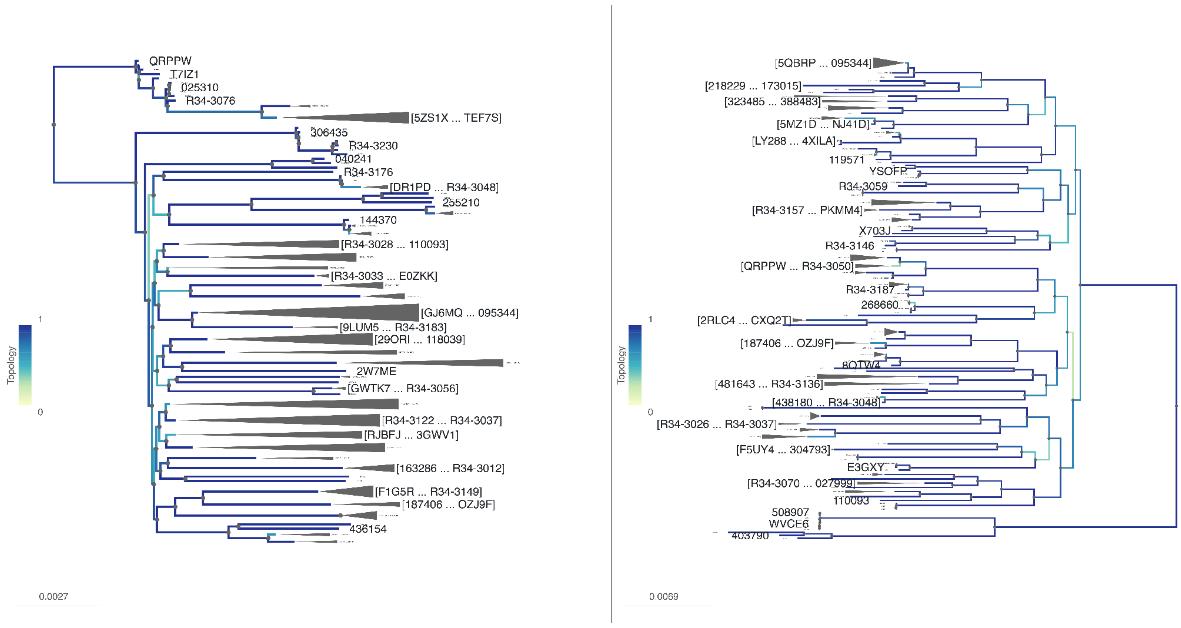


**Supplementary figure 2**: Comparison of original core-genome analysis and PopPIPE results in phylo.io, also available online:
https://beta.phylo.io/viewer/?session=766ab303a44c0ae882f543d7b163881a028f9e4e

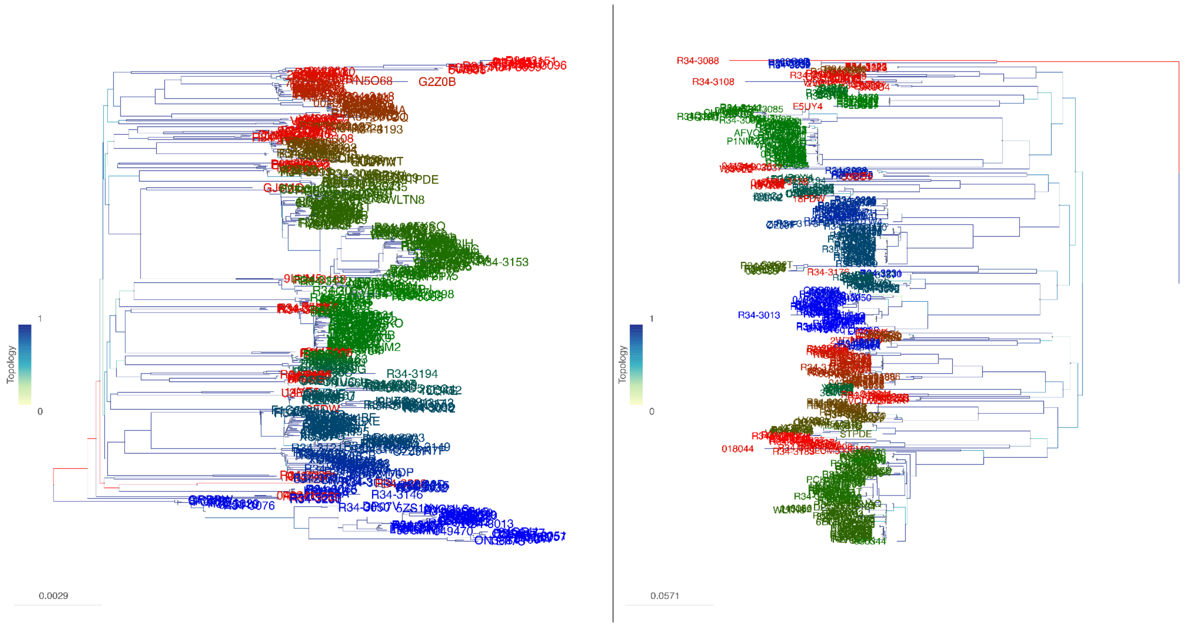


**Supplementary figure 3**: Comparison of original core-genome analysis and PopPIPE results in phylo.io including subcluster labels, also available online: [https://beta.phylo.io/viewer/?session=50677c9d8497401e949a2de9c64833feb528be9a](https://beta.phylo.io/viewer/?session=50677c9d8497401e949a2de9c64833feb528be9a)

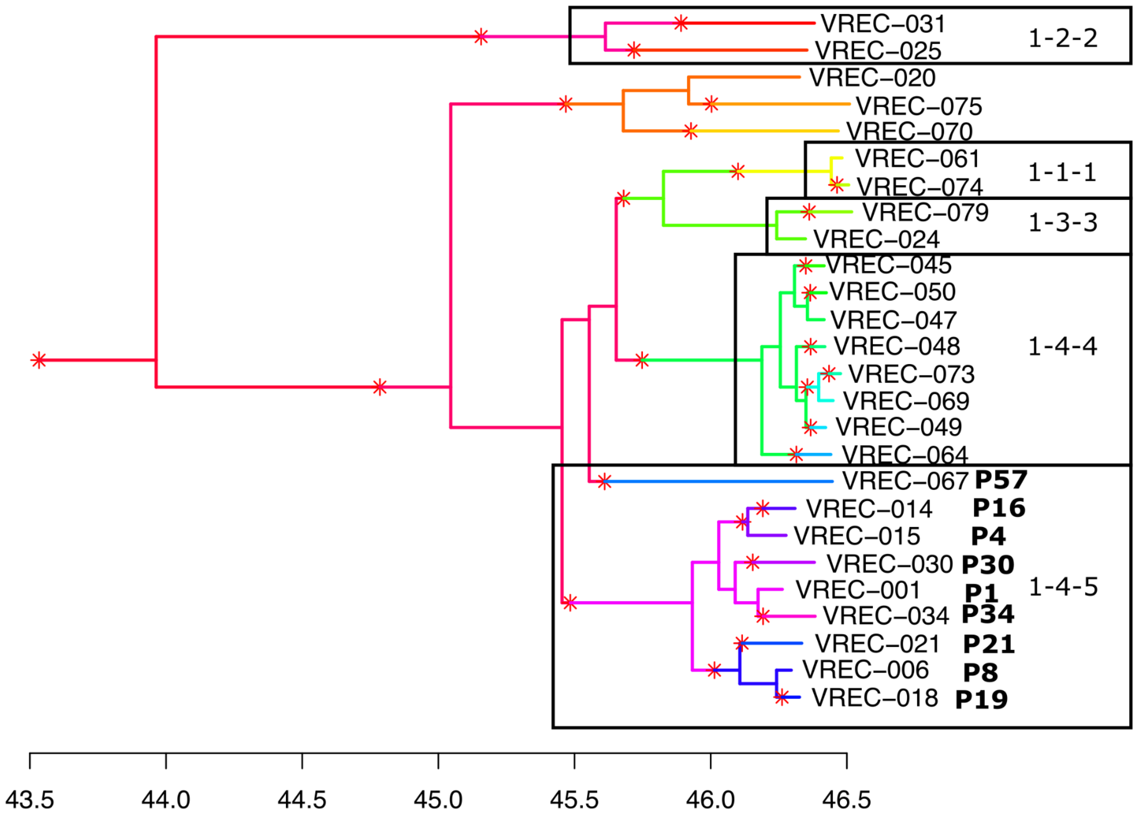


**Supplementary Figure 4:** TransPhylo transmission tree for PopPUNK cluster 1. PopPIPE clusters are indicated on the tree, and patient IDs have been added for cluster 1-4-5. Branches are coloured for each host, including inferred unsampled hosts, changes of colour on branches correspond to inferred transmission events from one host to another.
